# Development of the yeast *Saccharomyces* cerevisiae as a biosensor for the toxicity detection of toxic substances

**DOI:** 10.1101/2020.01.07.898106

**Authors:** Linlin Gong, Guang Yang, Bo Yang, Jihui Gu

**Affiliations:** Department of Medical Devices and Food, University of Shanghai for Science and Technology, Shanghai 200093, China

**Keywords:** yeast, culture conditions, chlorothalonil, sensitivity, biosensor, heavy metal

## Abstract

A whole-cell biosensor developed with yeast *Saccharomyces* cerevisia to detect the toxicity of chlorothalonil has been developed. This biosensor relied on the inhibition effect for metabolism by toxicants to provide detection and the degree of toxicity to yeast cells. In order to further improve the toxic sensitivity of yeast cells biosensor, the effect of the action time, the initial pH value of the medium and the temperature on inhibiting cell growth were investigated. Response surface regression analysis was conducted to obtain optimal culture conditions. Th effects of treated yeast morphology, ROS, DNA, caspase 3 activity were analyzed. This optimized yeast as a biosensor was used to detect chlorothalonil and heavy metals. The results are as follows: at optimal culture conditions, EC_50_ values of chlorothalonil to yeast biosensor determined at incubation time 4 h increased from 0.25 µg·mL^-1^ in the control to 0.006 µg·mL^-1^, which increased by 41.67 times. Compared with the control yeast cells, the morphology of optimized yeast cells were more transparent, with significantly increased intracellular vesicle rate and cell membrane permeability, intracelluar ROS increased siginificantly, DNA bands extracted was ladder, and caspase 3 activity was stimulated. The yeast biosensor had a high sensitivity to heavy metals. After analysis, many treated cells were apoptosis which was the main reason for the increasing sensitivity to detect harmful substances. It was found that the method provides a new idea for the detection of harmful substances in the environment.

Yeast cells biosensor could be used to detect harmful substances in the environment, sunch as chlorothalonil, heavy metals. Even through chemical analysis methods, such as ICP-MS and High Performance Liquid Choromatography (HPLC), have strengths in accuracy and limit of detection, it is impossible to evaluate the cytotoxicity and the biological effect of waste water by chemical result alone, and it is also expensive, prolix and complicated. However, the yeast cell biosensor is easy to operate, is sensitive to various toxicants, comparable to the other totxicity detection methods, is cheap in cost, and has. Therefore, the method which used yeast cells as biosensor will have great potential in the detection of the cytotoxicity of waste water in the future.

## 1. Instructions

Chlorothalonil (2,4,5,6-tetrachloroisophthalonitrile) is one of the most commonly used non-absorbable broad-spectrum fungicides, which is widely used in fruits, vegetables, grains and other crops[1]. Due to the widespread use of chlorothalonil, it is frequently reported that chlorothalonil remains in the atmosphere, water, soil and other environmental media[2-3]. According to the relevant toxicology experiments, chlorothalonil has accumulation effect in animals. If it exceeds a certain range, it will cause many serious chronic diseases. In especially, chlorothalonil has a certain connection with cancer[4]. Thus, chlorothalonil has been included in the list of chemical carcinogens in the United States[5].

At present, the main method to detect chlorothalonil is chemical analysis, but it does not conform to the twelve principles of green chemistry. They are usually expensive, with using a large number of toxic and environmentally harmful solvents. These solvents consume a lot of resources and produce harmful and toxic wastes [6]. Therefore, it is necessary to develop a reliable technology that can meet the requirements of green analytical chemistry, and the microbial inhibition method has the advantages of green environmental protection, low cost, simple and fast operation, and multifunctional detection combining analytical concentration and toxicity.

Biosensor is one of the important biological methods to detect harmful substances in the environment. In the early stage of biosensor development, fish, mouse algae, water flea and other individuals were mainly used. Using paramecium to detect the residual pesticide oxole pesticide in sewage [7]. Bee larvae have been used to detect chlorothalonil residues in water [8], however, due to the long growth cycle of individuals, it is not conducive to rapid detection requirements. With the establishment of cell-based system to evaluate the toxicity of harmful substances, at present, it is mainly used to detect the toxicity of harmful substances by using the characteristics of microbial physiological reaction, its own luminous characteristics or cell membrane. It has been studied to use *Saccharomyces* cerevisiae cells to determine the fungicide nitrogen disulfide, organic phosphorus pesticide and imidacloprid in waste water [9-10]. At present, the microbiological method mostly were utilizing yeast *Saccharomyces* cerevisia to detect estrogen in milk [11], and uses *Vibrio Fischer, E.coli*, pale bacteria to detect new harmful substances and heavy metals in waste water [12-14]. Although compared with the use of individual animals for toxicity detection, the microbiological method is low-cost, short-time and fast, it also has some disadvantages, such as the luminous bacteria must work in a certain salt solution and the number of cells would affect the optical signal and the detection error. In addition, it is believed that microorganisms with similar structure to higher organisms can more effectively respond to the toxicity of harmful substances. Yeast is a single cell eukaryote with a size of 10-20 μm. Because of its advantages of easy culture and rapid proliferation, yeast cell is regarded as an ideal model biomaterial in modern biological research [15], such as the acute toxicity of endocrine disruptors, sewage sediments, heavy metals and other chemicals [16-18].

The purpose of this study is to develop and optimize the yeast cell for the determination of chlorothalonil. In this experiment, yeast *Saccharomyces* cerevisia were utilized as a biosensor to detect the toxicity of chlorothalonil. The bioassay method has three important advantages: first, it can significantly improve the semi inhibition concentration of chlorothalonil; second, it can carry out toxicity test for different heavy metals; third, it has high accuracy and accuracy. Within a certain range of harmful substance concentration, the yeast cell OD_600_ is inversely proportional to the harmful substance concentration, and the yeast cell activity is weakened by changing the yeast cell culture conditions, so as to improve high concentration of chlorothalonil in yeast cells. Compared with the untreated yeast cells, the damaged yeast OD_600_ would change significantly. The results showed that the decrease of yeast OD_600_ under toxic and non-toxic conditions were directly proportional to the concentration of harmful substances, which provided the inhibition index. In addition, the toxicity of heavy metal Cu^2+^, Cd^2+^, Pb^2+^, Hg^2+^, Cd^6+^ in simulated waste water by yeast cell biosesor was introduced.

## 2. Materials and methods

### 2.1. Materials

High activity dry yeast was purchased from angel yeast company. The yeast cell were grown in YPD medium (1% yeast extract, 2% Bacto-peptone, 2% glucose, pH5.6) at 35°C with constant shaking (140 rpm), under aerobic condition.

Chlorothalonil was purchased from commerical sources and was prepared by dissolving 100 µg·mL^-1^ in water, which was then added to the medium at the concentrations indicated. The heavy metals used were ultrapure salts (CuSO_4_·5H_2_O, PbCl_2_, CdCl_2_, HgSO_4_ and K_2_Cr_2_O_7_). All chemical were of analytical grade. The other reagents were of reagent grade and purchased from commerical sources.

### 2.2. Yeast culture

0.1 g of active dry yeast powder was placed in 100 mL sterile water for 20 min at 35 °C. The activated yeast were grown overnight in 100 mL of YPD medium at 35 °C. The cell were harvested, centrifuged and washed, and then suspended in in a 100 mL fresh liquid medium. Then the OD_600_ value of the yeast liquid culture medium was measured by Ultraviolet visible spectrophotometry (UV-VIS, L5S, China) every two hours and finish after 24 h.

Yeast cells were harvested by centrifuging 5 mL aliquots of culture at 3,000 rpm 5 min, removing the supernatant and washing the resulting cell pellets three times in 0.9% (w/v) NaCl solution. Finally the cells were resuspended in saline for use [13].

### 2.3. Growth assays

To evaluate the effect of chlorothalonil on the growth of the yeasts, cells were suspended in 10 mL YPD medium to give an OD of 0.4 at 600 nm, and incubated time 4 h in various concentrations of chlorothalonil (0 µg ·mL^-1^ - 0.4 µg ·mL^-1^). Lately The cell growth was monitored turbidometry at 600 nm. Each treatment was carried out in triplicate. In addition, the cell inhibition rate (IR)was calculated as following:x

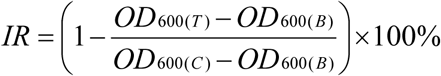

IR : cell inhibition rate(%); OD_600(T)_ : the OD_600_ value of yeast cells in YPD medium which containing chlorothalonil; OD_600(B)_ : the OD_600_ value of yeast cells in YPD medium is 0.4; OD_600(C)_ : the OD_600_ value of yeast cells in YPD medium.

### 2.4. The influence of culture conditions on yeast cell sensitivity to chlorothalonil

According to the preliminary experimental results, the action time, pH and temperature were taken as three single factors, and five levels were set as shown in table 1. The yeast OD_600_ was determined respectively. Each treatment was carried out in triplicate.

**Table 1.** Five level CCRD for three independent variables of the culture conditions of yeast.

| Single factor | Symbol | Level |  |  |  |  |
| --- | --- | --- | --- | --- | --- | --- |
| Action time ( h ) | A | 2 | 3 | 4 | 5 | 6 |
| The initial pH value of medium | B | 3 | 4 | 5 | 6 | 7 |
| Temperature( °C ) | C | 20 | 30 | 40 | 50 | 60 |

### 2.5. Sensitivity of yeast to chlorothalonil was analyzed by response surface method

The response surface test was designed based on the results of the above single-factor test. The level of design factors is shown in table 2.

**Table 2.** Factor level coding table.

| Level | Factor |  |  |
| --- | --- | --- | --- |
|  | A | B | C |
| -1 | 3 | 3 | 30 |
| 0 | 4 | 4 | 40 |
| 1 | 5 | 5 | 50 |

### 2.6. The sensitivity of optimized yeast to chlorothalonil

The yeast cells treated under the optimal culture conditions were suspended in YPD medium to give an OD of 0.4 at 600 nm, and incubated 4 h in various concentrations of chlorothalonil (0 µg · mL^-1^ to 0.012 µg · mL^-1^). Each treatment was carried out in triplicate, and the OD_600_ and the cell IR after 4 h exposure was calculated.

### 2.7. The influence of optimized conditions on yeast cell

The yeast cells treated under optimal culture conditions, were centrifuged at 3000 rpm 5 min, and the precipitate was washed twice with aseptic PBS, while the untreated cells as control.

#### 2.7.1. Morphology

The morphological changes of the yeast cells (1 × 10^5^) were analyzed using light microscope(NE610,China) at 1 000× and scanning electron microscope (SEM) at 50 000×.

#### 2.7.2. Cell membrane integrity

Cell membrane integrity was determined using lactate dehydrogenase (LDH) release assay kit. The cells were inoculated on the 96-well plate. Each culture hole was divided into four groups: the cell-free culture hole, the untreated control group, the untreated maximum enzyme activity control hole for subsequent cleavage, and the treated cell hole. The untreated cells were performed with 10% (W/V) reagent that releases LDH and used as a control group for maximum enzyme activity. LDH detection working fluid was added to each well and incubated in dark for 30min. The samples were read at 490 nm in a microplate reader (Readmax 1900,China).

#### 2.7.3. DHE staining

The reactive oxygen species (ROS) of yeast cells were measured by ROS assay kit that includes dihydroethidium (DHE). The yeast cells were treated with 5 μg·ml^-1^ DHE probe. Images of cells were captured using 40 × objectives under fluoresent microscope. The accumulation of ROS in the cells was determined according to the intensity of fluorescence intensity.

#### 2.7.4. DNA fragmentation

The yeast DNA was extracted from Ezup column yeast genomic DNA extraction kit and placed in 1.5% agarose gel electrophoresis apparatus on 120 V for 1.5 h. DNA fragmentation was detected by gel imager. The cells were divided into three group: the untreated control group, the heat death yeast cell group, and the treated yeast cell group.

#### 2.7.5. Capase 3 enzymatic activity

The protease activity of caspases 3 in yeast cells was performed utilizing a colorimetric assay kit. The resulting cells (2 × 10^6^) was lysed with 100 μL cold lysis buffer and inculbated on ice for 0.5 h. The resulting cell lysate was centrifuged at 10000 rpm for 10min and the suspernatant was collected. The activity of caspase 3 was measured immediately. The samples were read at 405 nm in a microplate reader.

### 2.8. Chemeical analysis and bioassays of the heavy metal waste water samples by yeast cells biosensor

Repeat test method 2.6 to detect heavy metal Cu^2+^, Pb^2+^, Hg^2+^, Cr^6+^ and Cd^2+^. Meanwhile, the waste water samples were filtered a 0.22 µm pore-size membrane filter, acidified by nitric acid before the measurement of heavy metals by Inductively Coupled Plasma Optical Emission Spectrometer (Agilent ICP-OES 5110, America). The test methods were carried out according to standard protocol(Method of determination for 26 elements (copper, nickel, lead, zinc, cadmium, chrome etc.) content in the sludge from industrial waste liquid treatment. GB/T 36690-2018).

### 2.9. Statistical analysis

Data analysis was performed using software Origin 8、SPSS 13.0 and MATLAB 2016. LSD method was used for comparison between the treatment group and the control group, and *P*<0.05 was defined as the difference. “*, **” respectively means *P<0.05, P<0.01*.

## 3. Results and disscussion

### 3.1. The growth curve of yeast

**Fig. 1.**
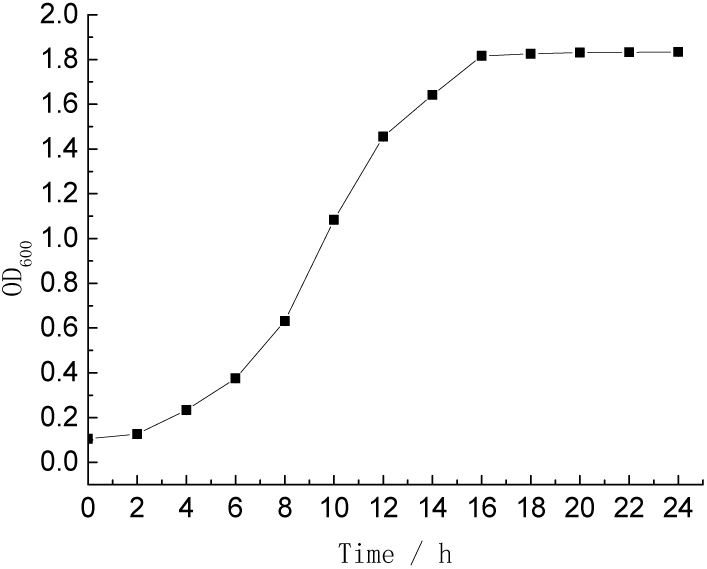
Growth curve of yeast in YPD medium.

Fig.1 indicates the growth curve of the yeast. As can be seen from it, 0∼6 h was the lag phase of yeast, 6∼16 h was the logarithmic phase of yeast, and especially 10 h was the middle timing of the logarithmic phase. After 14 h, yeast entered stationary phase and the growth curve tended to be stable. At 10 h, yeast cells had a strong ability to proliferate. Scariot et al also choose to culture yeast cells in this phase when studying the toxicity of dithianon to yeast cells[22]..Moreover, Wang et al show that the Psychrobacter sp. at the middle of logarithmic phase or transition from logarithmic to stationary phase would have a higher sensitivity to toxicants beasuse the phase would a strong growth rate and a higher metabolism. Therefore, yeast at 10 h was selected for subsequent experiments.

### 3.2. The effect of chlorothalonil on yeast proliferation

The toxicity of chlorothalonil on the proliferation of yeast strains, cells were grown in YPD medium, containing 0 to 0.4 µg ·mL^-1^. The cell growth was measured by monitoring the optical density at 600 nm. Meanwhile, according to test method 2.6, cell inhibition rates were also calculated. In figure 2, as for the cell growth curve, when the concentration of chlorothalonil was within the range of 0.4 µg · mL^-1^, the OD_600_ value of the yeast solution decreased with the increase of the concentration of chlorothalonil, indicating that the intensity of chlorothalonil inhibiting the proliferation of yeast increased with the increase of the concentration. When the concentration of chlorothalonil was 0.2∼0.3 µg · mL^-1^, the OD_600_ of yeast solution decreased the fastest. When the concentration of chlorothalonil was 0.4 µg · mL^-1^, the OD_600_ value of yeast solution did not change significantly compared with the initial concentration of bacteria solution, indicating that chlorothalonil almost completely inhibited the proliferation of yeast. In addition, cell inhibition rate increased with increasing concentration of chlorothalonil. Therefore, chlorothalonil has toxicity on yeast growth and inhibition rate, and present dosage dependence relations. The EC_50_ value, which represents the concentration of chlorothalonil that cause 50% inhibition of the growth rate, is about 0.25 µg ·mL^-1^. 0.4 µg ·mL^-1^ of chlorothalonil exerted the maximum inhibitory effect on yeast cell. 0.3 µg · mL^-1^ of chlorothalonil produced 87%, which is close to 100%(0.4 µg · mL^-1^). Considering the cost, the addition amount of chlorothalonil was chosen as 0.3 µg ·mL^-1^ in subsequent experiments.

**Fig. 2.**
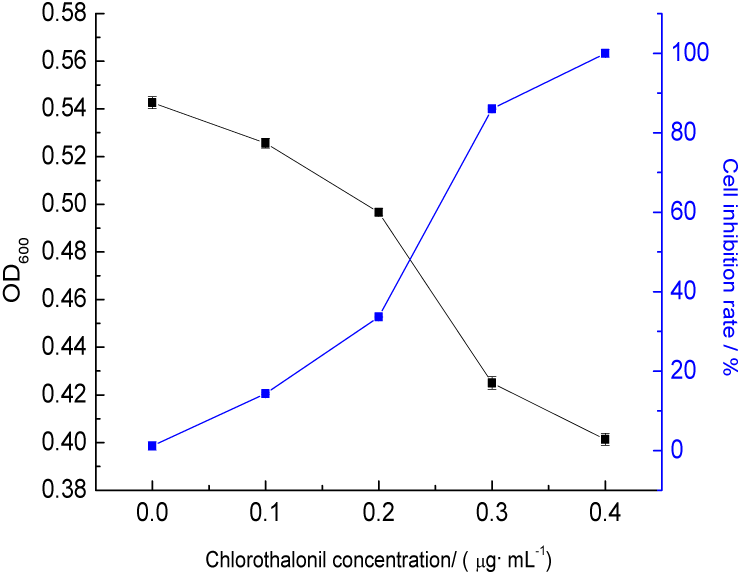
The effect of chlorothalonil on the growth of yeast.

### 3.3. Response surface analysis of yeast detection sensitivity

According to the results of the above single-factor experiment, the response surface optimization experiment was designed, and the results were shown in Table 3.

**Table 3.** Experimental results of response surface method.

| Run | A | B | C | Decrease of OD<sub>600</sub> value |
| --- | --- | --- | --- | --- |
| 1 | -1 | -1 | 0 | 0.195 |
| 2 | 1 | -1 | 0 | 0.192 |
| 3 | -1 | 1 | 0 | 0.196 |
| 4 | 1 | 1 | 0 | 0.181 |
| 5 | -1 | 0 | -1 | 0.163 |
| 6 | 1 | 0 | -1 | 0.152 |
| 7 | -1 | 0 | 1 | 0.168 |
| 8 | 1 | 0 | 1 | 0.168 |
| 9 | 0 | -1 | -1 | 0.156 |
| 10 | 0 | 1 | -1 | 0.155 |
| 11 | 0 | -1 | 1 | 0.165 |
| 12 | 0 | 1 | 1 | 0.159 |
| 13 | 0 | 0 | 0 | 0.214 |
| 14 | 0 | 0 | 0 | 0.215 |
| 15 | 0 | 0 | 0 | 0.212 |
| 16 | 0 | 0 | 0 | 0.214 |
| 17 | 0 | 0 | 0 | 0.213 |

Design Expert 8.05 software was used to regression fit the response surface test results, and the quadratic multiple regression model equation was obtained: Y= - 0.8308 + 0.0718a + 0.1243b +0.0331c - 0.003 ab + 0.0003 ac-0.0001bc - 0.009a^2^ - 0.0133 b^2^ - 4.1550 c^2^, and the results were shown in Table 4. The model P < 0.01, the missing item P=0.1104 > 0.05, the correlation coefficient R^2^=99.78%. The results show that the model fits well. CV was 0.93%, indicating that the test results were feasible. Therefore, the test results could be analyzed and predicted.

**Table 4.** Analysis of variance model of regression equation.

| Source | Sum of squares | DF | mean square | F value | P(Prob>F) | significant |
| --- | --- | --- | --- | --- | --- | --- |
| Model | 0.0030 | 9 | 0.0003 | 353.16 | < 0.0001 | ** |
| A | 0.0000 | 1 | 0.0000 | 35.98 | 0.0005 | ** |
| B | 0.0000 | 1 | 0.0000 | 12.37 | 0.0098 | ** |
| C | 0.0001 | 1 | 0.0001 | 49.46 | 0.0002 | ** |
| AB | 0.0000 | 1 | 0.0000 | 12.32 | 0.0099 | ** |
| AC | 0.0000 | 1 | 0.0000 | 10.35 | 0.0147 | * |
| BC | 0.0000 | 1 | 0.0000 | 2.14 | 0.1870 |  |
| A<sup>2</sup> | 0.0005 | 1 | 0.0005 | 124.65 | < 0.0001 | ** |
| B<sup>2</sup> | 0.0008 | 1 | 0.0008 | 254.94 | < 0.0001 | ** |
| C<sup>2</sup> | 0.0012 | 1 | 0.0012 | 2488.19 | < 0.0001 | ** |
| Residual | 0.0000 | 7 | 0.0000 |  |  |  |
| Lack of fit | 0.0000 | 3 | 0.0000 | 3.91 | 0.1104 |  |
| Pure error | 0.0000 | 4 | 0.0000 |  |  |  |
| Total | 0.0030 | 16 |  |  |  |  |

In Table 4, A, B, C, AB, A^2^, B^2^ and C^2^ show significant differences, and AC interaction show significant differences. The response surface analysis of the regression model was conducted with design-expert software, and the results were shown in Fig 3, Fig 4, and Fig 5. The Optimization module of the software was used to optimize the culture conditions. When the predicted action time was 3.95 h, the initial pH value was 3.97, and the culture temperature was 40.42°C, OD_600_ decreased by 0.2141. According to the actual test conditions, the temperature was adjusted to 40.4°C, yeast was cultured under the above optimal conditions, and three parallel experiments were conducted. The actual OD_600_ decrease average value was 0.2139. The experimental results were close to the predicted value, which proved that the regression model could better predict the influence of culture conditions on the sensitivity of yeast to detect chlorothalonil.

**Fig. 3.**
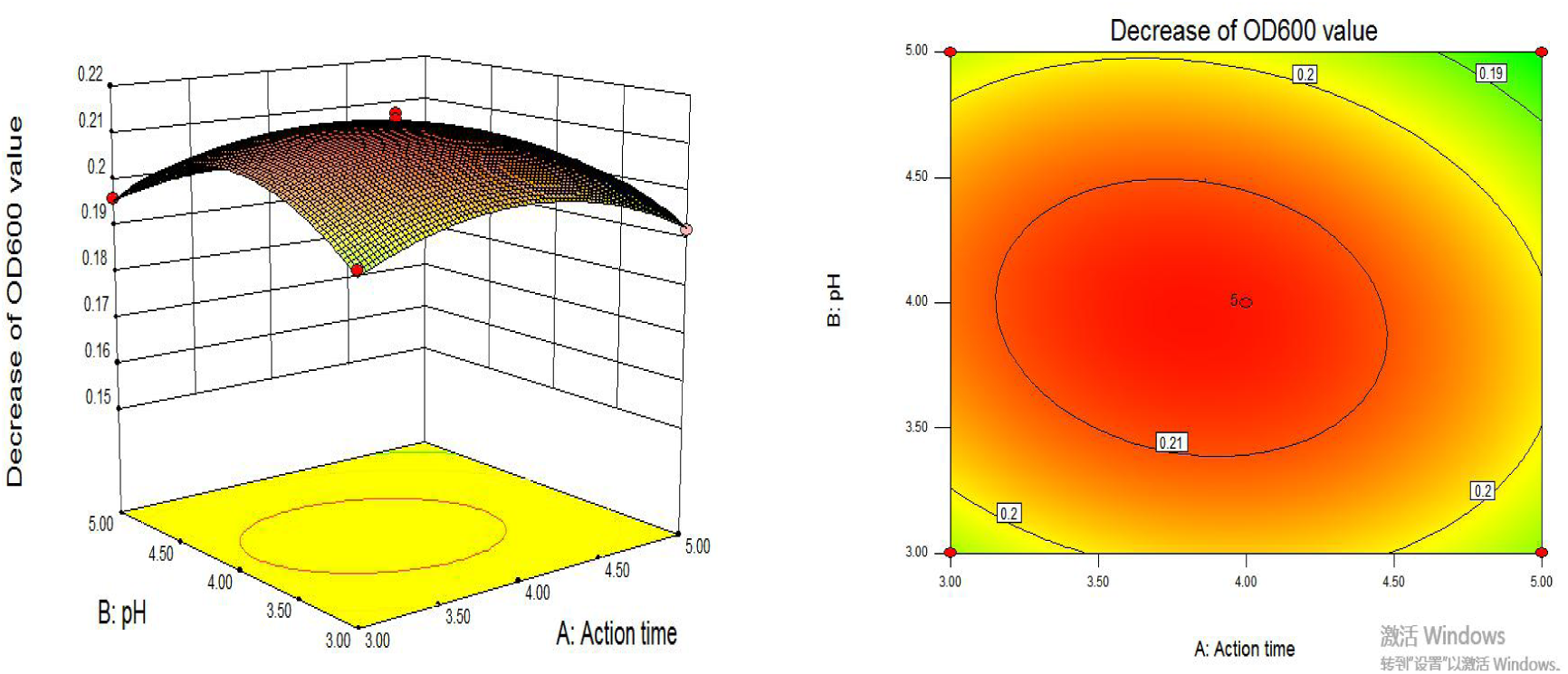
The effect of action time and pH on sensitivity of yeast detection.

**Fig. 4.**
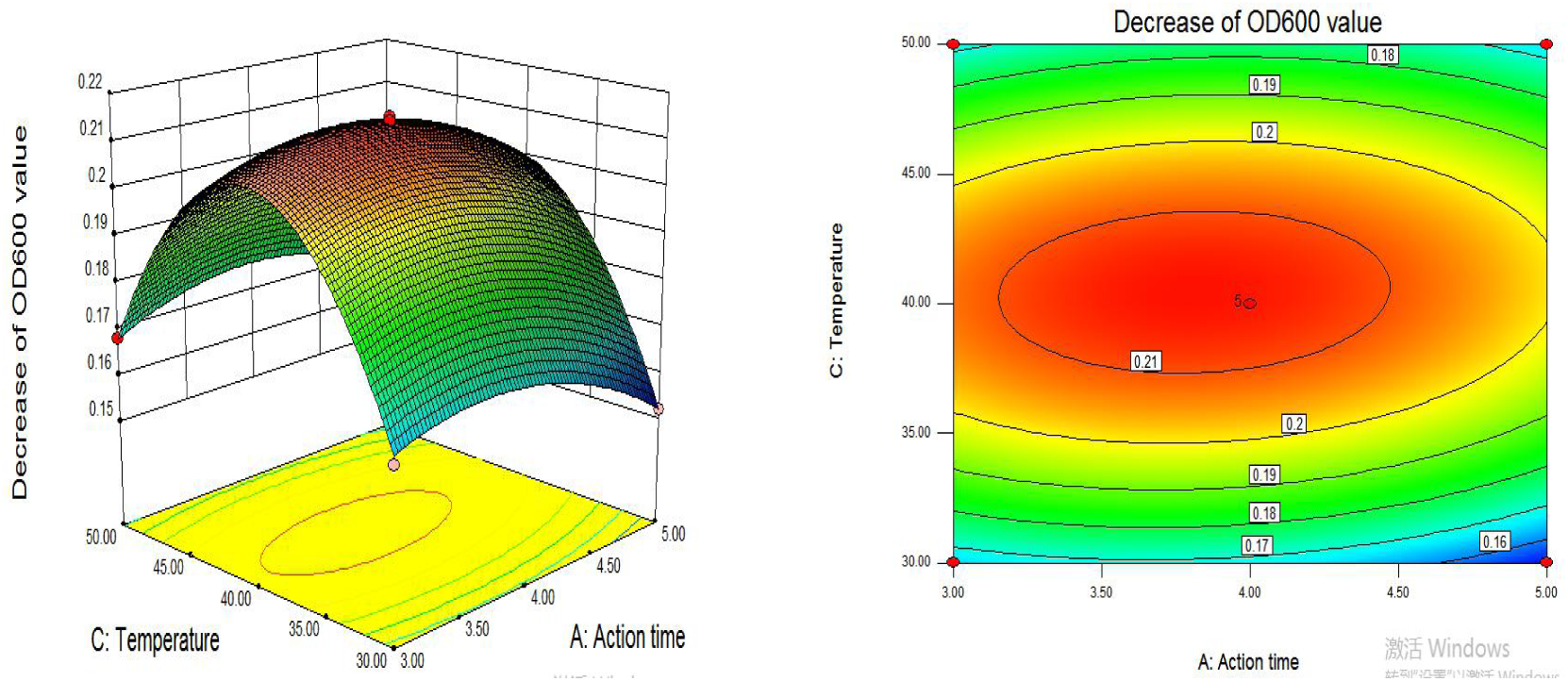
The effect of time and temperature on sensitivity of yeast detection.

**Fig. 5.**
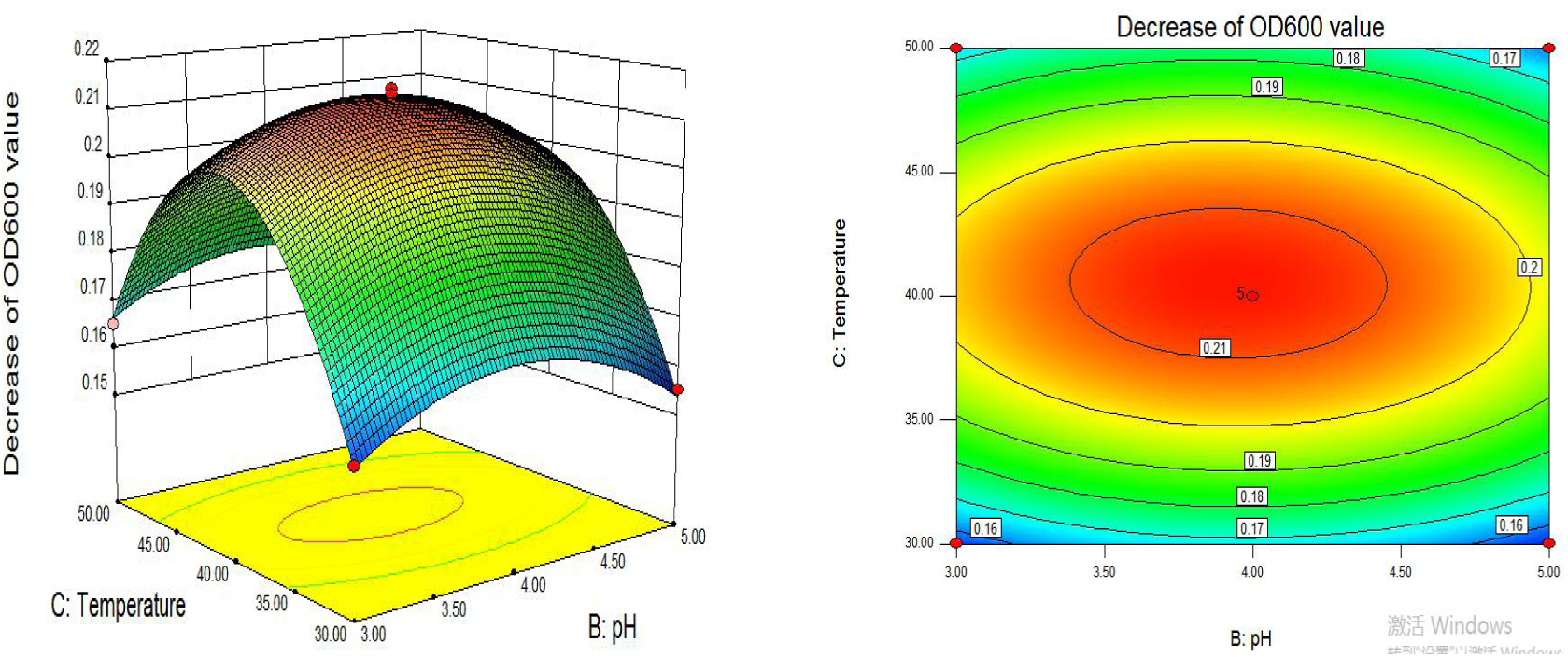
The effects of pH and temperature on sensitivity of yeast detection.

### 3.4. The sensitivity of yeast detection under optimal response surface conditions

The optimal experimental conditions obtained from the response surface were as following: the action time was 3.95 h, the initial pH of the medium was 3.97, and the culture temperature was 40.4 °C. Yeast was cultured and its sensitivity to the detection of chlorothalonil was measured, which were as shown in Fig 6. The concentration of chlorothalonil was within the range of 0 µg· mL^-1^∼0.004 µg · mL^-1^, and the OD_600_ value began to decrease. Chlorothalonil had an inhibitory effect on the proliferation of yeast. When the concentration of chlorothalonil was between 0.004 µg · mL^-1^ - 0.008 µg · mL^-1^, the OD_600_ value decreased significantly, and the intensity of chlorothalonil’s effect on yeast cell proliferation gradually increased. The EC_50_ value represents the concentration of chlorothalonil that caused 50% inhibition of the growth rate. Compared the EC_50_ value of the untreated yeast cell, it is easy found that the EC_50_ value of yeast on chlorothalonil decreased from 0.25 µg · mL^-1^ to 0.006 µg · mL^-1^ (Fig. 2 and Fig. 6), which indicated that the sensitivity of yeast to chlorothalonil could be improved after culture.

**Fig. 6.**
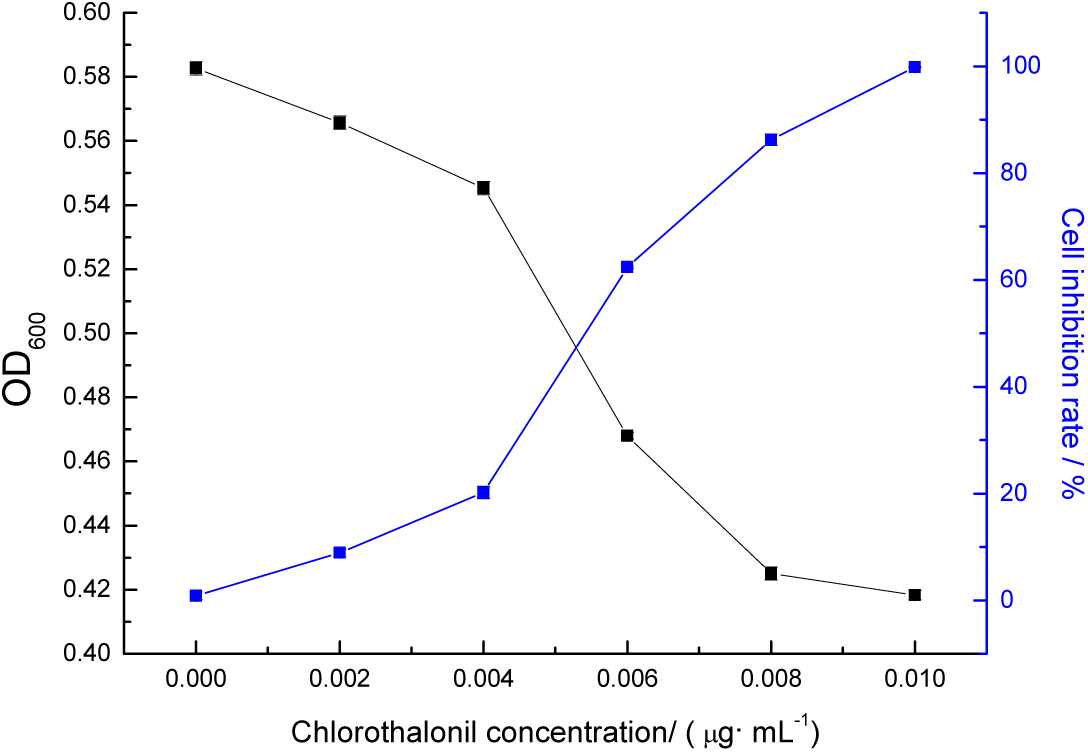
The effects of chlorothalonil on yeast growth.

### 3.5 The influence of optimized conditions on yeast cell

#### 3.5.1. Morphology

In order to explore the influence of optimized culture conditions on the internal structure of yeast cells, the optimized yeast cells were observed with a optical microscope and SEM (Fig. 7). In control group, the internal structure of control cells was relatively complete, fat particles accumulated in large quantities, and cells hardly had vesicles. But under the optimal culture condition, the cell fat granule was decreased in the internal structure, and the percentage of cavitation was significantly increased (Fig. 8). In addition, the overall transparency of the cell was significantly improved.

**Fig. 7.**
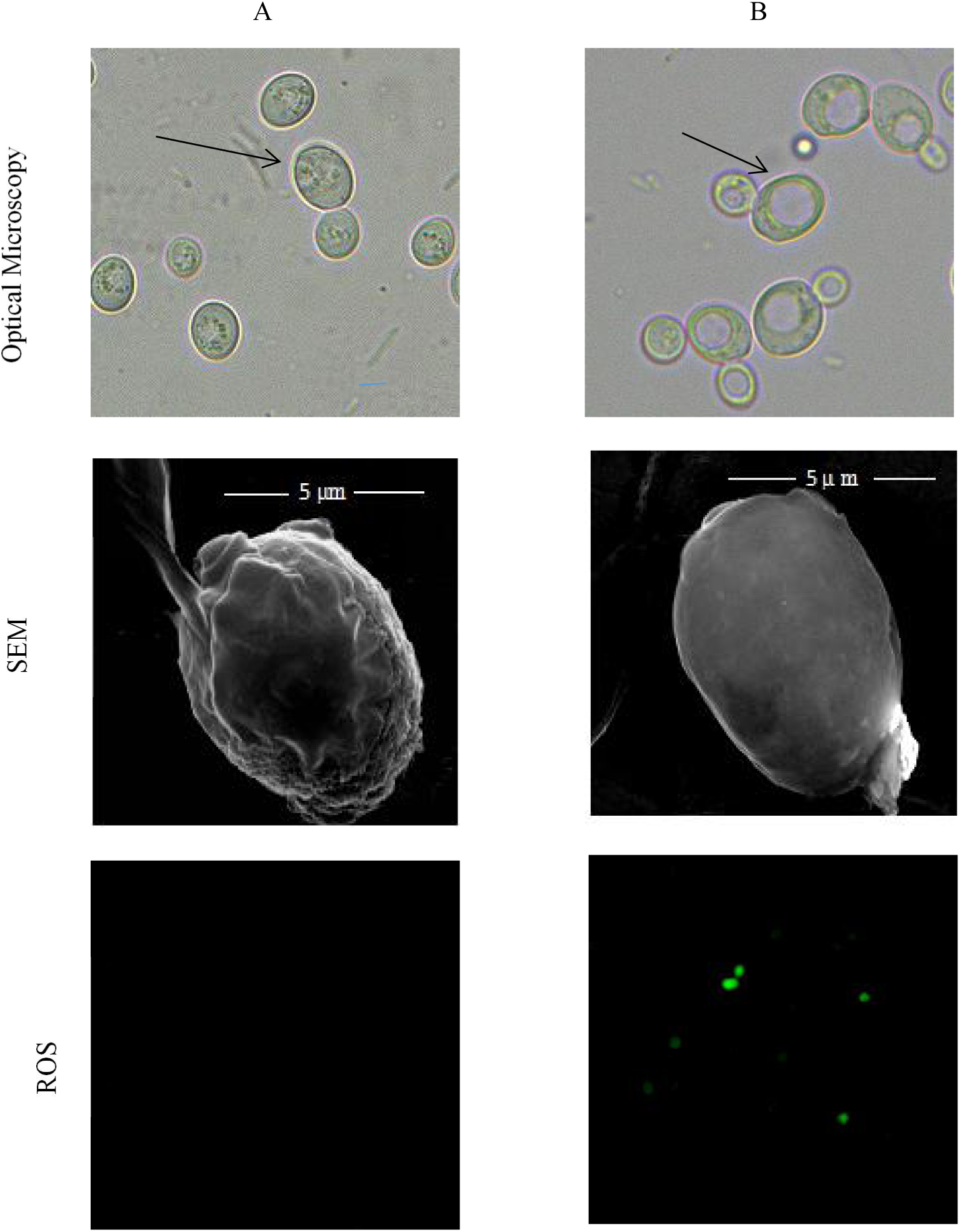
Effects of optimized culture conditions on the internal structure of yeast cells. The cells were grown under aerobic condition. Aliquots were taken at the indicated time. A, yeast cells in YPD medium at 35°C;

**Fig. 8.**
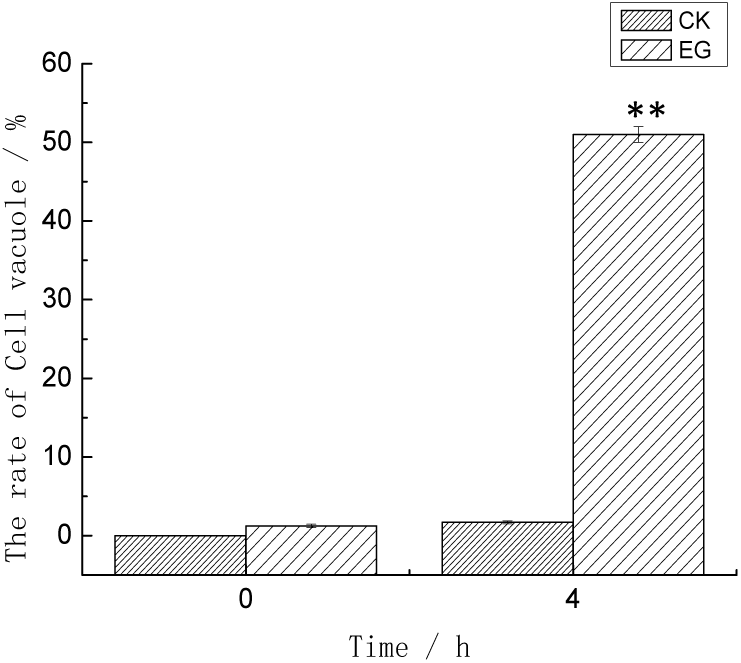
Effects of optimized culture conditions on the Internal Structure.

At the same time, the cells were also observed by SEM (Fig.7). In control group, the surface folds and villi of yeast cells were obvious. But in the optimum culture conditions of experimental group, the surface of yeast cells were smooth and no fold, and bacteria shape was from round to oval, for the reason that the yeast cells maybe morbid, and cell wall maybe thinner or more loose structure, and cell membrane permeability could increase, which leading that intracellular and extracellular harmful substances are more likely to enter the bacteria cause more damage, reduce the cell internal structure, and then cells become more fragile. The vesicle occurs the phenomenon with acetic acid in yeast cells consistent with the experimental results and internal microstructure morphology [22]. Therefore, it is useful that changing yeast culture conditions in a certain extent would improve the sensitivity of chlorothalonil yeast cells.

#### 3.5.2 Measure of ROS generation

In order to determine to the possible association of ROS in optimized conditions-induced cavitation, the ROS of yeast cells was measured to using fluorescent probe DHE in the present study. As shown in Fig. 7, in control, no observable fluorescence images were found. However, after yeast cells were treated under optimized culture conditions, some bright green fluorescence cells were easily observed, which is because intracelluar ROS level increased. When the ROS generation is lower than the threshold, it will cause apoptosis, and when the threshold is exceeded, it will cause death. In apoptosis the increased ROS would result in decreased mitochhondrial permeability. In apoptosis, the increased ROS would result in decreased mitochhondrial permeability, which indirectly leaded to increased capase 3 activity.

#### 3.5.3 DNA fragmentation

DNA ladder of yeast cells was a characteristic of apoptosis. In the present study, to further distinguish the treated yeast cells between apoptosis or necrosis. The yeast DNA was extracted and placed in 1.5% agarose gel electrophoresis apparatus on 120 V for 1.5 h. DNA fragmentation was detected by gel imager. As shown in Fig. 9, in control, DNA electrophoresis band consists of only one large molecular fragment, DNA electrophoresis bands of heat death cells were diffuse without visible, and DNA electrophoresis bands of treatment yeast cells was ladder, which is due to the action of endonuclease.

**Fig. 9.**
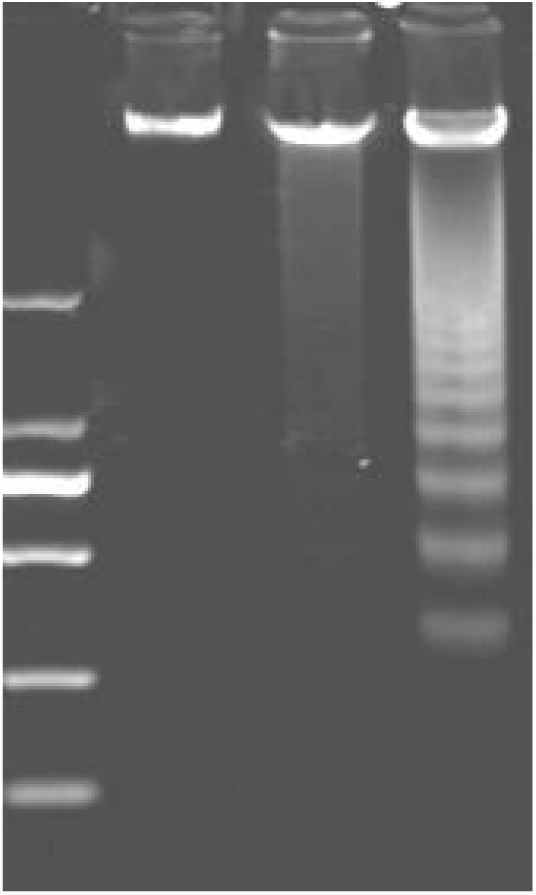
DNA electrophoresis band. 2000 bp DNA ladder (lane 1), DNA extracted from control group (lane 2), DNA extracted from heat death group (lane 3), DNA extracted from treated group (lane 4).

#### 3.5.4 Caspase 3 activity

To investigate the involvement of caspase signaling cascade in optimized conditions-induced apoptosis, yeast cells were treated under optimized conditions and caspase 3 enzymatic was determined. Compared to control groups, caspased 3 of the treatment was much higher(Table 5). In apoptosis, caspase would result in cavitation. The phenomenon was according to the results of the morphology.

**Table 5.** Caspase 3 activity unit of yeast cell.

| Group | A<sub>405</sub> | pNA/ μM | Caspase 3 activity unit /U |
| --- | --- | --- | --- |
| Control | 0.0419 | 0.901 | 0.901 |
| Treat | 0.088 | 42.43 | 42.43 |

Taken together, the results confirm that the treatment group of optimized conditions increased ROS generation, which stimulated caspase 3 activity with cavaitation and induced apopotosis with DNA ladder, which were also verified that the effective inhibition concentration of chlorothalonil against yeast increased from 0.25 µg· mL^-1^ to 0. 006 µg· mL^-1^, which increased by 41.67 times.

### 3.6 Toxicities of heavy metal to yeast cells of optimized conditions

In order to verify the optimization can be used as a harmful material detection sensitivity, the important heavy metal pollutants, such as Cu^2 +^, Cd^2 +^, Pb^2 +^, Hg^2 +^, Cd^6 +^ were respectively investigated under the optimal experimental conditions in present study. The responses of yeast cell based biosensor to the heavy metal ions were shown in Table 6. The EC_50_ results obtained were compared with those of published data by other methods (Table 6). As shown in Table 6, the EC_50_ values of toxicity assay based on yeast cells were comparable and even better than those of other tests.The results were similar to Wang et al.’s report on toxicity order of heavy metals based on *Psychrobacter* sp. biosensor. Analysis of the reason may be due to different methods. Those toxicity assays were used electricall and optical real-time response for toxicity[20]. As for Hg^2+^, it s much lower toxicity than other heavy metals was mainly due to its stronger membrane sequestration. Moreover, according to the 4 h-EC_50_ value of the five measured heavy metals to yeast cell biosensor, the toxicity was in order of Cd^2+^>Cu^2+^>Pb^2+^>Cr^6+^>Hg^2+^. The toxicity of Cd^2+^ is strong, because 0.5 µg · mL^-1^ of Cd^2+^ could inhibition 50% of growth of yeast cells. However, Cui et al report that Hg^2+^ is strong in the toxicity[23]. Therefore, there was some significant difference in the sensitivities of the various tests and methods, which reflected the fact that each bacterial species and test procedure had its own sensitivity pattern to toxicants.

As shown in Table 7 and Fig. 10. The OD_600_ value of the yeast solution decreased with increasing concentration of Cu^2 +^, Cd^2 +^, Pb^2 +^, Hg^2 +^, and Cd^6+^ and exhibited a significant correlation with the following concentration of Cu^2 +^(0 µg· mL^-1^ - 4 µg · mL^-1^), Cd^2 +^(0 µg · mL^-1^ - 0.08 µg · mL^-1^), Pb^2+^(0 µg · mL^-1^ - 4 µg · mL^-1^), Hg^2+^(0 µg · mL^-1^ - 8 µg · mL^-1^), and Cd^6+^(0 µg · mL^-1^ - 6 µg · mL^-1^). The calibration cures were shown in table 6, which x was toxicants’ concentration and y was absorbance. It is found that the method using yeast cell biosensor has a strong correlation in toxicity test for five heavy metals. Moreover, The ICP-OES methods were carried out according to standard protocol(GB/T 36690-2018). There was also significant correlation between the signal value (CPS) of internal standard elements and the concentrations of measured elements(data no shown). Even though the working curve of ICP-OES is high significant, the method can not detect the cytotoxicity[]. In this case, the yeast cell biosensor would make it a good method to detect cytotoxicity of heavy metals in waste water, such as Cu^2 +^, Cd^2 +^, Pb^2 +^, Hg^2 +^, and Cd^6+^.

**Table 7.** Comparison of EC_50_ value by toxicity assays to others.

| Toxicity assays | Toxicants, $\text{EC}_{50}$ ($\mu\text{g} \cdot \text{mL}^{-1}$) | | | | | References |
| --- | --- | --- | --- | --- | --- | --- |
| | $\text{Cu}^{2+}$ | $\text{Cd}^{2+}$ | $\text{Pb}^{2+}$ | $\text{Hg}^{2+}$ | $\text{Cr}^{6+}$ | |
| Yeast cells biosensor, incubation time 4 h | 2.1 | 0.5 | 2.8 | 3 | 2.5 | Present study |
| *Acinetobacter baylyi* Tox2 biosensor, incubation time 15 min | - | 43.67 | - | 0.12 | - | Cui et al., 2018 |
| M. Chula fish, exposure time 96 h | 0.68 | 39.83 | - | 0.14 | - | Cui et al., 2018 |
| *Psychrobacter* sp. biosensor, incubation 30 min | 2.6 | 47.3 | 110.1 | 0.8 | 14 | Wang et al., 2013 |
| FM-TOX,*E.coli*, incubation time 60 min | 3.7 | 7.8 | 20.4 | 2.0 | - | Catterall et al., 2010 |

**Fig. 10.**
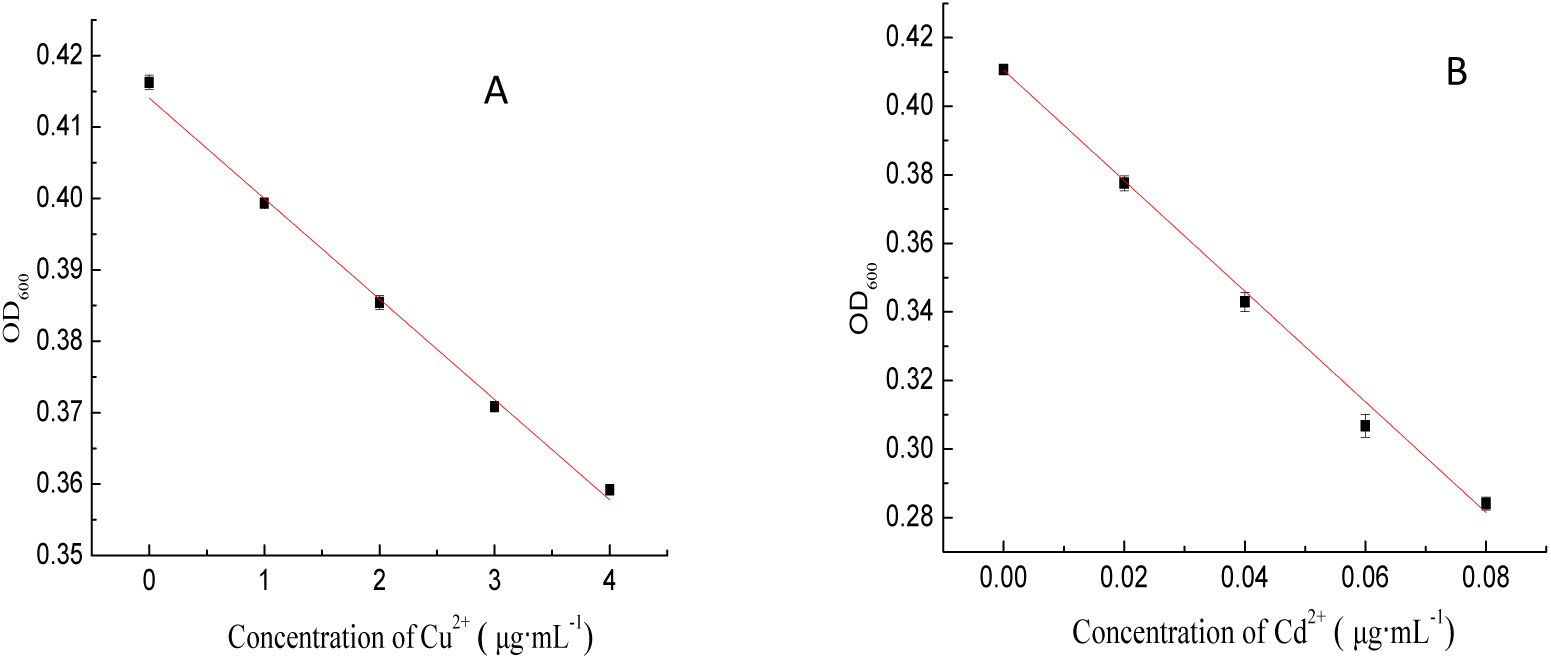
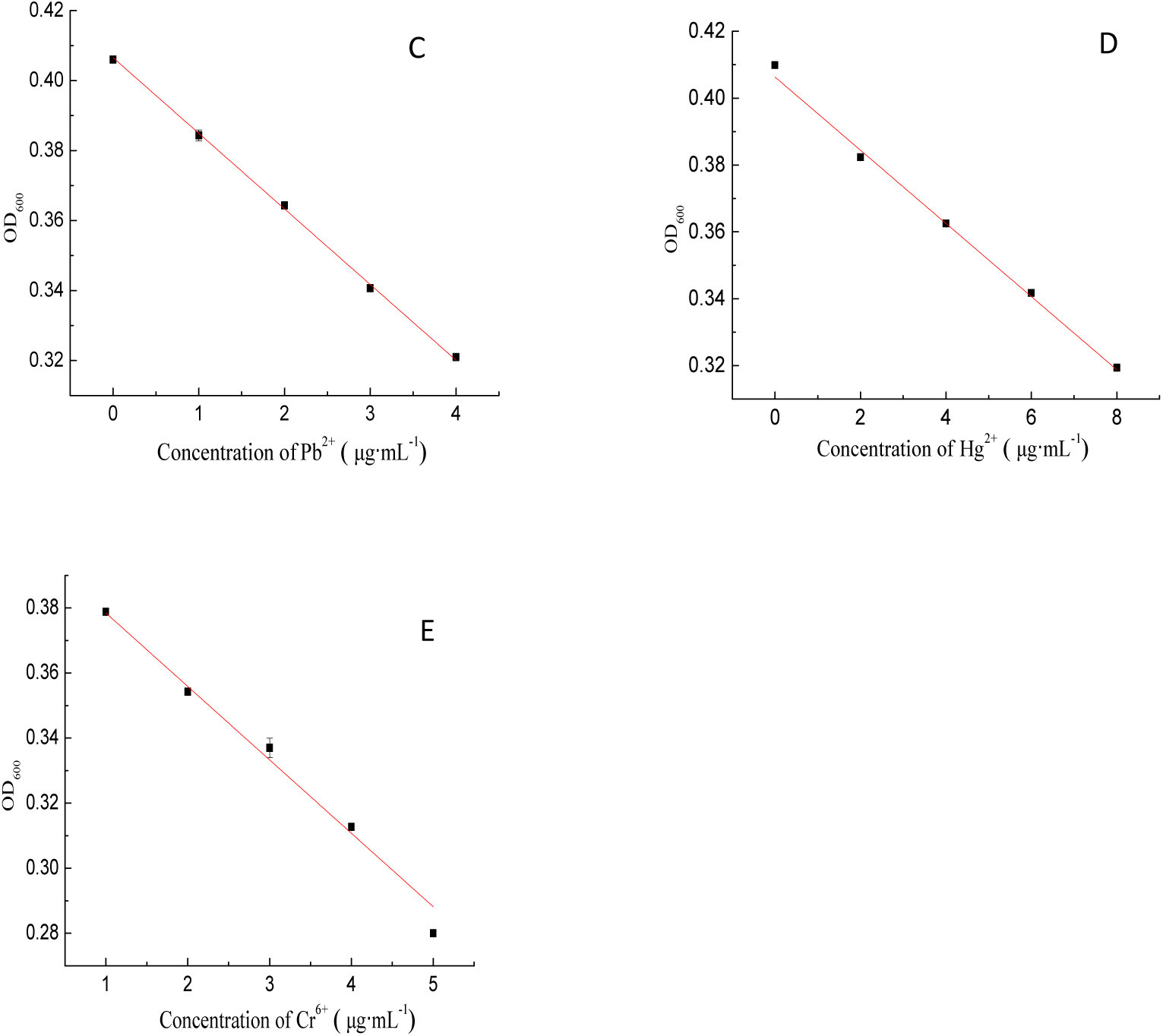
Regression of yeast cell biosensor under different concentrations of Cu^2+^(A), Cd^2+^(B), Pb^2+^(C), Hg^2+^(D), Cr^6+^(E).

The reproducibility of yeast cell biosensor was evaluated by six replicate assays for heavy metal ions Cu^2 +^, Cd^2 +^, Pb^2 +^, Hg^2 +^ and Cr^6 +^ inhibition on yeast cell during 4 h incubation. The relative standard deviations (R.S.D.) obtained and the rate of recovery obtained were shown in table 8. Wang et al evaluated the reproducibility of the *Psychrobacter* sp. biosensensor by five heavy metals, Cu^2+^, Cd^2+^, Zn^2+^, Cr^6+^, which would inhibited on *Psychrobacter* sp. during 30 min incubation. The R.S.D. were 6.5%, 8.2%, 7.0%,7.3%,2.5%, and 9.1%, respectively. However, all of the R.S.D. of yeast cell biosensor by five heavy metals, Cu^2+^, Cd^2+^, Zn^2+^, Cr^6+^, which would inhibited on *Psychrobacter* sp. During 4 h incubation, were 0.15%, 0.17%, 0.21%, 0.14% and 0.18%, respectively. It is easy fount that the R.S.D. are lower than the above. In addition, the recovery rate obtained were 101.11%, 103.86%, 100.10, 100.54%, 101.29%, respectively. Because the test data were in the range of 95%-105%, which meet the requirement of recovery rate. Thus, the yeast cells biosensor showed a good, reproducible behavior and could be used for reproducible measurements. Moreover, with a portable and compact design, the yeast cells biosensor is able to monitor toxicity in different location.

**Table 8.** The results of toxicities of heavy metal to yeast cell biosensor.

| Toxicants | Range<br>($\mu\text{g} \cdot \text{mL}^{-1}$) | Regression | $R^2$ | p | RSD<br>(%) | Recovery rate<br>(%) |
| --- | --- | --- | --- | --- | --- | --- |
| $\text{Cu}^{2+}$ | 0-4 | $y = -0.01394x + 0.4561$ | 0.9938 | <0.01 | 0.15 | 101.11 |
| $\text{Cd}^{2+}$ | 0-0.08 | $y = -1.499x + 0.410$ | 0.9980 | <0.01 | 0.17 | 103.86 |
| $\text{Pb}^{2+}$ | 0-4 | $y = -0.0216x + 0.40654$ | 0.9979 | <0.01 | 0.21 | 100.10 |
| $\text{Hg}^{2+}$ | 0-8 | $y = -0.0096x + 0.40378$ | 0.9960 | <0.05 | 0.14 | 100.54 |
| $\text{Cr}^{6+}$ | 0-5 | $y = -0.02527x + 0.4561$ | 0.9914 | <0.01 | 0.18 | 101.29 |

The reproducibility of yeast cell biosensor was evaluated by six replicate 355 assays for heavy metal ions Cu^2 +^, Cd^2 +^, Pb^2 +^, Hg^2 +^ and Cr^6 +^ inhibition on yeast cell during 4 h incubation. The relative standard deviations (R.S.D.) obtained and the rate of recovery obtained were shown in table 8. Wang et al evaluated the reproducibility of the Psychrobacter sp. biosensensor by five heavy metals, Cu^2+^, Cd^2+^, Zn^2+^, Cr^6+^, which would inhibited on Psychrobacter sp. during 30 min incubation. The R.S.D. were 6.5%, 8.2%, 7.0%,7.3%,2.5%, and 9.1%, respectively. However, all of the R.S.D. of yeast cell biosensor by five heavy metals, Cu^2+^, Cd^2+^, Zn^2+^, Cr^6+^, which would inhibited on Psychrobacter sp. During 4 h incubation, were 0.15%, 0.17%, 0.21%, 0.14% and 0.18%, respectively. It is easy fount that the R.S.D. are lower than the above. In addition, the recovery rate obtained were 101.11%, 103.86%, 100.10, 100.54%, 101.29%, respectively. Because the test data were in the range of 95%-105%, which meet the requirement of recovery rate. Thus, the yeast cells biosensor showed a good, reproducible behavior and could be used for reproducible measurements. Moreover, with a portable and compact design, the yeast cells biosensor is able to monitor toxicity in different location.

Even though chemical analysis methods, such as ICP-MS and High Performance Liquid Choromatography (HPLC), have strengths in accuracy and limit of detection, it is impossible to evaluate the cytotoxicity and the biological effect of waste water by chemical result alone, and it is also expensive, prolix and complicated. However, the yeast cell biosensor is easy to operate, is sensitive to various toxicants, comparable to the other totxicity detection methods, is cheap in cost, and has. Therefore, the method which used yeast cells as biosensor will have great potential in the detection of the cytotoxicity of waste water in the future.

## 4. Conclusion

In this experiment, the culture condition, such as the action time, medium initial pH, and temperature, could affect the yeast cells’ sensitivity of chlorothalonil. Response surface methodology was used to increase the effects of culture conditions on yeast cells’ sensitivity of chlorothalonil. There was the optimal culture conditions: the action time of 3.95 h, medium initial pH value of 3.97, culture temperature is 40.4 °C. In this case, the OD_600_ decreased to 0.2141, which is the maximum in all teats. The optimization of culture conditions for yeast detection sensitivity was validated, the actual OD_600_ dropped to 0.2139, which was close to the predicted value. Thus the response surface test results more reliable. Under the optimal culture conditions, the EC_50_ value which responses inhibitory concentration of chlorothalonil on yeast cell decreased from 0.25 µg · mL^-1^ to 0.006 µg · mL^-1^ is due to many treated cells were apoptosis, which indicated that the sensitivity of yeast to chlorothalonil was improved. The response surface test results were further verified by observing yeast morphology.

According to China’s national standard (food pesticide maximum residue limits (GB 2763-2016)), chlorothalonil maximum residue in agricultural products (called simply MRL) in 0.1 ∼ 70 mg/kg. Chlorothalonil has been included in the list of chemical carcinogens in the United States. Compared to the maximum residue limits of our country of chlorothalonil in food, the yeast cells’ sensitivity of chlorothalonil under the condition of the optimal cultivation was higher than that, which meets the detection limit of chlorothalonil residue in food. In addition, the optimal yeast cells would be used to detect heavy metals in waste water, evaluate the bioavailability and the biological effect of waste water, and the accuracy and reproducibility were good. Thus, the method of using yeast cells as biosensor to detect the residue of chlorothalonil in food provides a new idea for the detection of heavy metals in the environment.

## Acknowledgements

Authors acknowledge the authorities of Shanghai University of technology for providing the facilities to carry out this work.

## References

1 Guerreiro A D S, Rola R C, Rovani M T, et al. Antifouling biocides: Impairment of bivalve immune system by chlorothalonil[J]. Aquat Toxicol, 2017, 189:194–199.

2 Wu X, Yin Y, Wang S, et al. Accumulation of chlorothalonil and its metabolite, 4-hydroxychlorothalonil, in soil after repeated applications and its effects on soil microbial activities under greenhouse conditions[J]. Environmental Science and Pollution Research, 2014, 21(5): 3452∼3459.

3 Arinaitwe K, Kiremire B T, Muir D C G, et al. Legacy and currently used pesticides in the atmospheric environment of Lake Victoria, East Africa[J]. Science of the Total Environment, 2016, 543(PtA):9∼18.

4 EPA, 1999. Chlorothalonil ⟨ http://archive.epa.gov/pesticides/reregistration/web/pdf/0097fact.pdf ⟩. (Accessed on 14 December 2017).

5 FAO/WHO. Pesticide Residues in Food. FAO Plant Production and Protection. 1992, 116: 29–31.

6 Wieczerzak M, Namies Nik J, KudłAk B. Bioassays as one of the Green Chemistry tools for assessing environmental quality: A review[J]. Environment International, 2016, 94:341∼361.

7 Hussain M M, Amanchi N R, Solanki V R, et al. Low cost microbioassay test for assessing cytopathological and physiological responses of ciliate model Paramecium caudatum to carbofuran pesticide[J]. Pesticide Biochemistry and Physiology, 2008, 90(1):66–70.

8 Pingli D, Jack C J, Mortensen A N, et al. The impacts of chlorothalonil and diflubenzuron on Apis mellifera L. larvae reared in vitro[J]. Ecotoxicology and Environmental Safety, 2018, 164:283–288.

9 Scariot F J, Luciane J, Delamare A P L, et al. Necrotic cell death induced by dithianon on, Saccharomyces cerevisiae[J]. Pesticide Biochemistry and Physiology, 2018:149:137-142.

10 Liang B, Lu X, Wang G, et al. Functional cell-surface display of acetylcholinesterase for spectrophotometric sensing organophosphate pesticide[J]. Sensors and Actuators B: Chemical, 2018, 279: 483–489.

11 Westlund P, Yargeau V. Investigation of the presence and endocrine activities of pesticides found in waste water effluent using yeast∼based bioassays[J]. Science of the Total Environment, 2017, 607-608:744.

12 StypulA-TreBas S, Minta M, Radko L, et al. Application of the yeast∼based reporter gene bioassay for the assessment of estrogenic activity in cow’s milk from Poland[J]. Environmental Toxicology and Pharmacology, 2015, 40(3):876–885.

13 Znag Y, Ren T, He J, et al. Acute heavy metal toxicity test based on bacteriahydrogel[J]. Colloids and Surfaces A, 2019, 563:318–323.

14 Kusumahastutia K.A. D, Sihtmäeb M, Kapitanov Illia V, et al. Toxicity profiling of 24 L∼phenylalanine derived ionic liquids based on pyridinium, imidazolium and cholinium cations and varying alkyl chains using rapid screening Vibrio fischeri bioassay[J]. Ecotoxicology and Environmental Safety, 2019, 172:556–565.

15 Braconi D, Bernardini G, Santucci A. *Saccharomyces cerevisiae* as a model in ecotoxicological studies: A post-genomics perspective[J]. Journal of proteomics, 2015, 137:19–34.

16 Hettwer K, JäHne M, Frost K, et al. Validation of, Arxula, Yeast Estrogen Screen assay for detection of estrogenic activity in water samples: Results of an international interlaboratory study[J]. Science of The Total Environment, 2018, 621:612–625.

17 Leskinen P, Hilscherova K, Sidlova T, et al. Detecting AhR ligands in sediments using bioluminescent reporter yeast[J]. Biosensors & Bioelectronics, 2008, 23(12):1850–1855.

18 Rumlova L, Dolezalova J. A new biological test utilising the yeast *Saccharomyces* cerevisiae for the rapid detection of toxic substances in water[J]. Environmental Toxicology and Pharmacology, 2012, 33(3):0–464.

19 Scariot Fernando J, Jahn Luciane, Delamare Ana Paula L, et al. Necrotic cell death induced by dithianon on Saccharomyces cerevisiae[J]. Pesticide biochemistry and physiology, 2018, 149.

20 Xuejiang W, Mian L, Xin W, Zhen W, et al. P -benzoquinone-mediated amperometric biosensor developed with *Psychrobacter* sp. for toxicity testing of heavy metals[J]. Biosensors and Bioelectronics, 2013, 41.

21 Qi Xiang, L, Panpan,L, Peng,H, et al. Dual-signal-biosensor based on luminescent bacteria biofilm for real-time online alert of Cu(II) shock.[J]. Biosensors & bioelectronics, 2019, 142.

22 Yachen D. Analysis of programmed cell death induced by acetic in Saccharomyces cerevisiae based on omics technology [D]. Zhejiang University, 2015.

23 Zhisong C, Luan X, Huichao J, et al. Application of a bacterial whole cell biosensor for the rapid detection of cytotoxicity in heavy metal contaminated seawater.[J]. Chemosphere, 2018, 200.

